# Immunoinformatics approach for a novel multi-epitope vaccine construct against spike protein of human coronaviruses

**DOI:** 10.1101/2021.05.02.442313

**Authors:** Avinash Kumar, Ekta Rathi, Suvarna G Kini

**Affiliations:** Department of Pharmaceutical Chemistry, Manipal College of Pharmaceutical Sciences, Manipal Academy of Higher Education, Manipal, Karnataka, India

**Keywords:** Covid-19, Coronaviruses, Cheminformatics, Immunoinformatic, SARS-CoV-2, Vaccine

## Abstract

Spike (S) proteins are an attractive target as it mediates the binding of the SARS-CoV-2 to the host through ACE-2 receptors. We hypothesize that the screening of S protein sequences of all the HCoVs would result in the identification of potential multi-epitope vaccine candidates capable of conferring immunity against various HCoVs. In the present study, several machine learning-based *in-silico* tools were employed to design a broad-spectrum multi-epitope vaccine candidate against S protein of human coronaviruses. To the best of our knowledge, it is one of the first study, where multiple B-cell epitopes and T-cell epitopes (CTL and HTL) were predicted from the S protein sequences of all seven known HCoVs and linked together with an adjuvant to construct a potential broad-spectrum vaccine candidate. Secondary and tertiary structures were predicted, validated and the refined 3D-model was docked with an immune receptor. The vaccine candidate was evaluated for antigenicity, allergenicity, solubility, and its ability to achieve high-level expression in bacterial hosts. Finally, the immune simulation was carried out to evaluate the immune response after three vaccine doses. The designed vaccine is antigenic (with or without the adjuvant), non-allergenic, binds well with TLR-3 receptor and might elicit a diverse and strong immune response.

## 1. Introduction

The increased incidence of infectious diseases, especially of zoonotic origin, might become an existential threat to human species. In 2018, WHO (world health organization) had come out with a blueprint of priority disease list which identified viruses like Nipah, Ebola and MERS (middle east respiratory syndrome) posing international biosafety threats(“Blueprint,” n.d.). It also waved a red flag on the pandemic risk potential of the infections caused by known zoonotic pathogens and collectively classified these threats as “disease X”(“Blueprint,” n.d.). The fears of 2018 came true in the early months of 2020 when Covid-19 emerged as the most devastating pandemic the world has ever seen. Around 3.15 million people have lost life with numbers rising daily and more than 150 million people reported positive to the SARS-CoV-2 (Severe Acute Respiratory Syndrome Coronavirus 2) virus responsible for Covid-19(“Home -Johns Hopkins Coronavirus Resource Center,” n.d.). A recent study suggests that coronaviruses (CoVs) and bats have co-evolved for millions of years but CoVs seldom jumped across to infect humans(Joffrin et al., 2020). They were considered as a comparatively harmless pathogen causing mild respiratory illness to humans(Song et al., 2019). But in the last two decades after the incidences of SARS infection in 2003 and MERS infection in 2013, the understanding regarding CoVs have changed. SARS-CoV-2 has been reported to target pneumocytes which cause severe respiratory distress characterized by bronchiolitis, bronchitis and pneumonia(Abraham Peele, Srihansa, Krupanidhi, Ayyagari, & Venkateswarulu, 2020). The current pandemic of Covid-19 is unprecedented in many ways and its impact on the economies around the world has been devastating. The scientific community has thrown all its resources to come up with either a drug to treat Covid-19 or a vaccine as a prophylactic measure with limited success.

SARS-CoV-2 like SARS and MERS belongs to the *Coronaviridae* family. The presence of spikes on the outer surface of CoVs has a resemblance of the crown and hence the name corona (in Latin corona means crown)(Pandey et al., 2020). SARS-CoV-2 is an enveloped, positive-sense, single-stranded RNA beta-coronavirus with 26-32 thousand base pairs(de Wilde, Snijder, Kikkert, & van Hemert, 2018). Its genome encodes both structural like E (envelop) protein, N (nucleocapsid) protein, M (membrane) protein, S (spike) protein and non-structural proteins (nsp) like proteases(Mousavizadeh & Ghasemi, 2020). Out of all the four structural proteins, S protein plays a prominent role in the binding of the SARS-CoV-2 to the host target receptors, especially through ACE-2 (angiotensin-converting enzyme 2) receptor(Wrapp et al., 2020; Zhou et al., 2020). Cleavage of S protein by furin or other proteases has been reported in many of the avian and mammalian CoVs. The cleavage of S results into S1 which bears the receptor-binding domain (RBD) and S2 responsible for fusion(Wrapp et al., 2020). Many authors have suggested that RBD of S protein might be an interesting target for potential therapies against SARS-CoV-2(Kadioglu1, Saeed1, Johannes Greten2, & Efferth1, n.d.; Panda et al., 2020; Prajapat et al., 2020). Scientists have achieved significant success in deciphering the key structural and non-structural proteins of the virus. Wrapp et al. have reported a Cryo-EM structure of S protein of SRAS-CoV-2 in the prefusion conformation(Wrapp et al., 2019). Zhang et al. have reported the x-ray crystal structure of the unliganded SARS-CoV-2 M^pro^ and its complex with an α-ketoamide inhibitor(L. Zhang et al., 2020). The researchers have employed various cheminformatics tools to target these proteins and develop potential drugs for Covid-19. Recently, several authors have reported SARS-CoV-2 M^pro^ inhibitors by using molecular modeling methods like virtual screening, molecular docking, molecular dynamics simulations, fast pulling of ligand (FPL), and free energy perturbation (FEP)(Alexpandi, De Mesquita, Pandian, & Ravi, 2020; Ferraz, Gomes, S Novaes, & Goulart Trossini, 2020; Gentile et al., 2020; Ngo, Quynh Anh Pham, Thi Le, Pham, & Vu, 2020; Olubiyi, Olagunju, Keutmann, Loschwitz, & Strodel, 2020). Hagar et al. have investigated some antiviral N-heterocycles as potential Covid-19 drug by using molecular docking and DFT calculations(Wrapp et al., 2019).

But apart from the search of drugs for Covid-19, researchers have tried to develop a vaccine against it. The trust in vaccine development comes primarily from the experience of successful mitigation of epidemics caused by coronaviruses in farm animals by vaccines(F. Li & Du, 2019; Song et al., 2019). Vaccines against human coronavirus which are under development comprise of DNA and RNA vaccines, vector-based vaccine, live attenuated virus itself or protein subunit-based vaccines(Corbett et al., 2020; Mateus et al., 2020; Sahin et al., 2020; van Doremalen et al., 2020). Epitope-based vaccine approach has been employed to combat several diseases like solid tumor(Knutson, Schiffman, & Disis, 2001), malaria(R. Wang et al., 2004) and multiple sclerosis(Bourdette et al., 2005). Recently, many authors have reported epitope-based vaccine candidates for the mitigation of Covid-19(Abdelmageed et al., 2020; Abraham Peele et al., 2020; Enayatkhani et al., 2020; Mukherjee, Tworowski, Detroja, Mukherjee, & Frenkel-Morgenstern, 2020; Naz, Shahid, Butt, Awan, & Ali, 2020; Oany, Emran, & Jyoti, 2014; Sarkar, Ullah, Tuz, Afrin, & Araf, 2020). Yang et al. have combined immunoinformatic methods with the deep learning algorithm to directly predict potential vaccine subunits from the S protein sequence of SARS-CoV-2 and constructed a multi-epitope vaccine for SARS-CoV-2 virus(Z. Yang, Bogdan, & Nazarian, 2020). Similarly, Singh et al. Have designed multi-epitope vaccine peptides by targeting four different structural proteins (Spike, Envelop, Membrane and Nucleocapsid) of SARS-CoV-2(A. Singh, Thakur, Sharma, & Chandra, 2020). Li et al. have also employed several immunoinformatic tools to design a multi-epitope vaccine by targeting three different structural proteins of SARS-CoV-2(Lin et al., 2020).

When we looked into the recent literature, we found most of the designed multi-epitope vaccine were against the S protein of SARS-CoV-2 virus(Dar et al., 2020; Kar et al., 2020; Naz, Shahid, Butt, Awan, Ali, et al., 2020; Sanami et al., 2020; H. Singh, Jakhar, & Sehrawat, 2020; Tahir ul Qamar et al., 2020). Hence, in the present study, we also decided to target the S protein but not of only SARS-CoV-2 but all the seven known strains of human coronaviruses (HCoVs) i.e. both alpha (HCoV-NL63 and HCoV-229E) and beta coronaviruses (HCoV-OC43, HCoV-HKU1, MERS, SARS-CoV and SARS-CoV-2). As S protein is a common structural protein among the HCoVs, hence, we have taken up the S protein sequence to design a multi-epitope vaccine candidate which might confer immunity against all these HCoVs. Our hypothesis of identifying an epitope capable of providing immunity against various CoVs is also supported by the recent work of Mateus et al., where they have identified cross-reactive T-cell epitopes of SARS-CoV-2 in unexposed humans. They have concluded that pre-existing immunity could be derived from the exposure to other HCoVs like HCoV-OC43, HCoV-229E, HCoV-NL63, or HCoV-HKU1, which cause the common cold(Mateus et al., 2020). Yang et al have reported that a vaccine candidate targeted towards the RBD of the S protein of SARS-CoV-2 led to the induction of protective immunity(J. Yang et al., 2020). To the best of our knowledge, it is the first such work where S proteins of all the seven known HCoVs were screened to design broad-spectrum multiepitope vaccine candidate which might confer immunity from all the seven HCoVs. We have taken extra care to select T-cell epitopes and hence, the predicted T-cell epitopes from different web servers were further filtered based on epitope conservancy analysis and molecular docking analysis with HLA-A*24:02 protein. Further, the best docked epitope-protein complex was also analyzed using MD simulations study for 50 ns. Thus, for the final vaccine construct, not only top-ranked predicted T-cell epitopes were considered but also based on conservation across different HCoV strains and peptide docking analysis. Finally, our designed vaccine candidate was analyzed for its potential to induce immune response through immune simulation studies and when compared to other recent publications, our designed vaccine candidate induced a higher level of immunoglobins (IgM + IgG), immunocomplexes and interleukins (IL-2). The vaccine candidate showed a minimum conservancy of 73.97% across the seven HCoVs and hence, it might provide protection against all these HCoVs. A flowchart summarizing the methodology and tools employed for the current work has been depicted in Fig. 1.

**Fig. 1.**
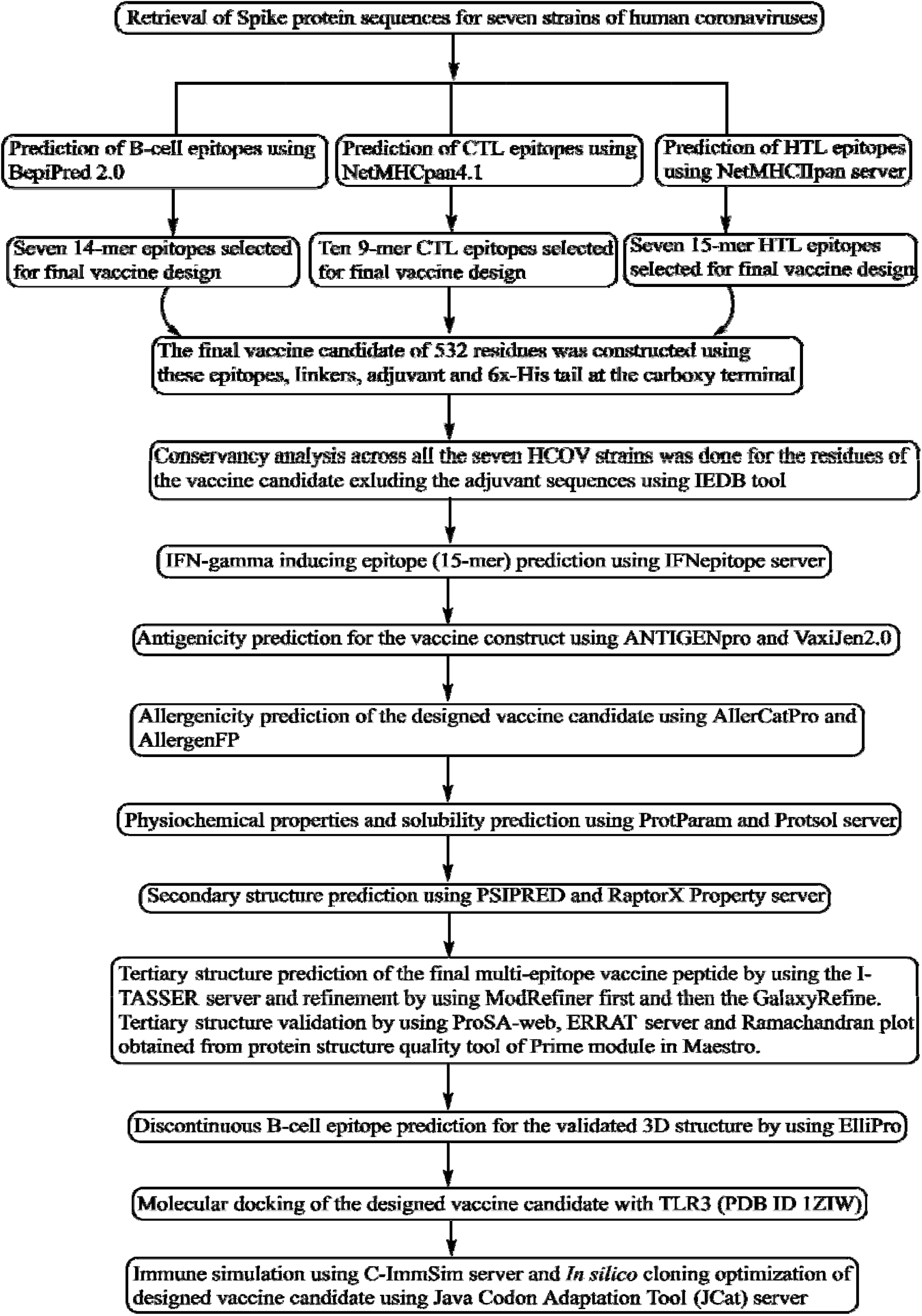
A flowchart summarizing the methodology and tools employed for the current work.

### 2. Materials and Methods

### 2.1 Experimental

All the immunoinformatic exercises were carried out by using free web-based tools. For cheminformatics experiments like molecular docking and molecular dynamics simulations were carried out by using Maestro (Release 2018-3) which is a commercial small-molecule drug discovery suite from Schrodinger Inc. (USA) on an HP computer with Linux Ubuntu 18.04.1 LTS operating system. It had Intel Haswell graphics card, 8GB RAM and Intel Core i3 processor.

### 2.2 Retrieval of S protein sequences

For the present study, we have taken the sequences of S protein from seven strains of coronaviruses (CoVs): (i) NL63 (ii) 229E (iii) OC43 (iv) HKU1 (v) MERS (vi) SARS-CoV and (vii) SARS-CoV-2. The sequences were retrieved in the FASTA format from the UniProtKB database(Apweiler et al., n.d.). The sequences of S protein from all the CoVs have been shown in Table 1.

**Table 1.**
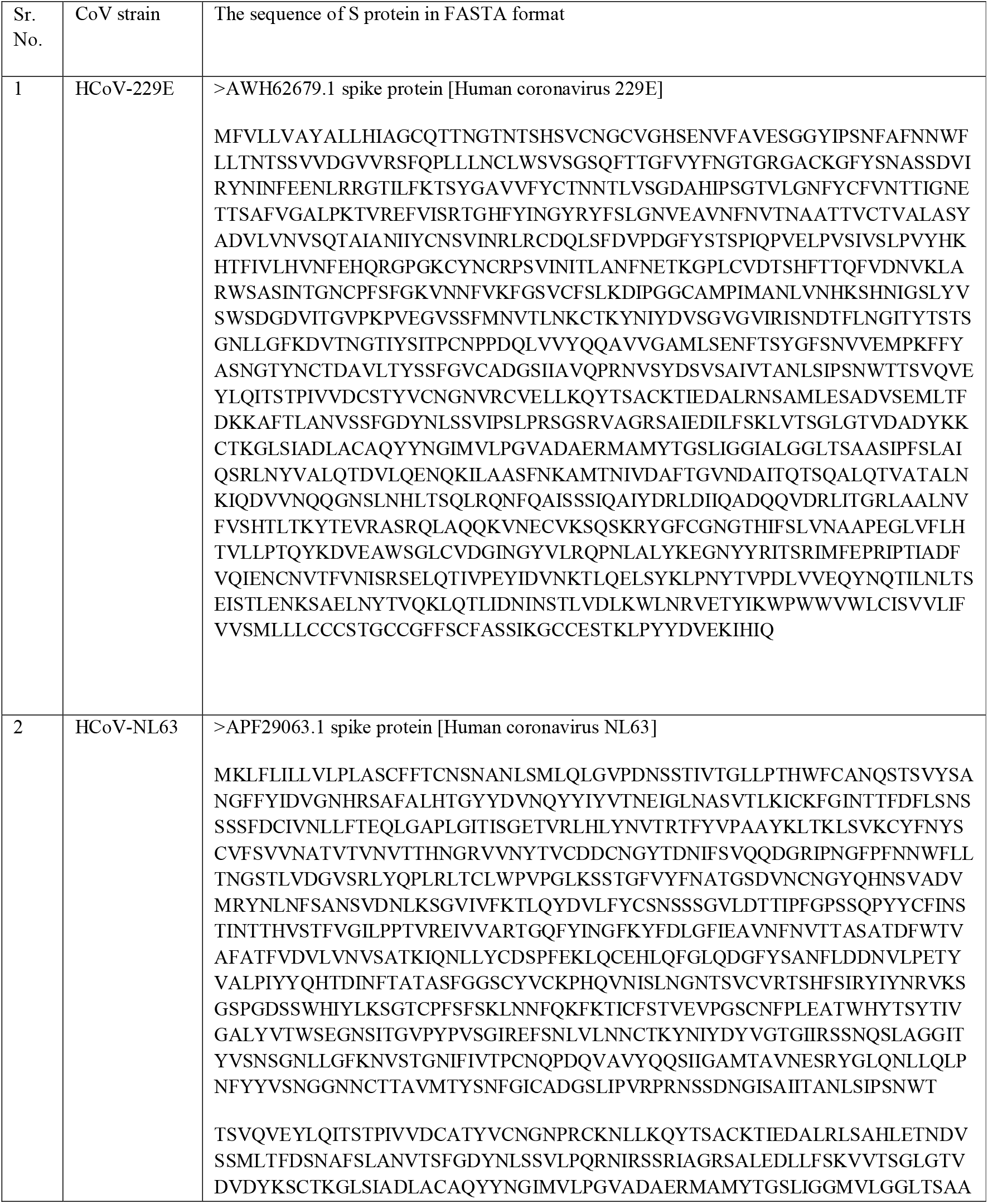
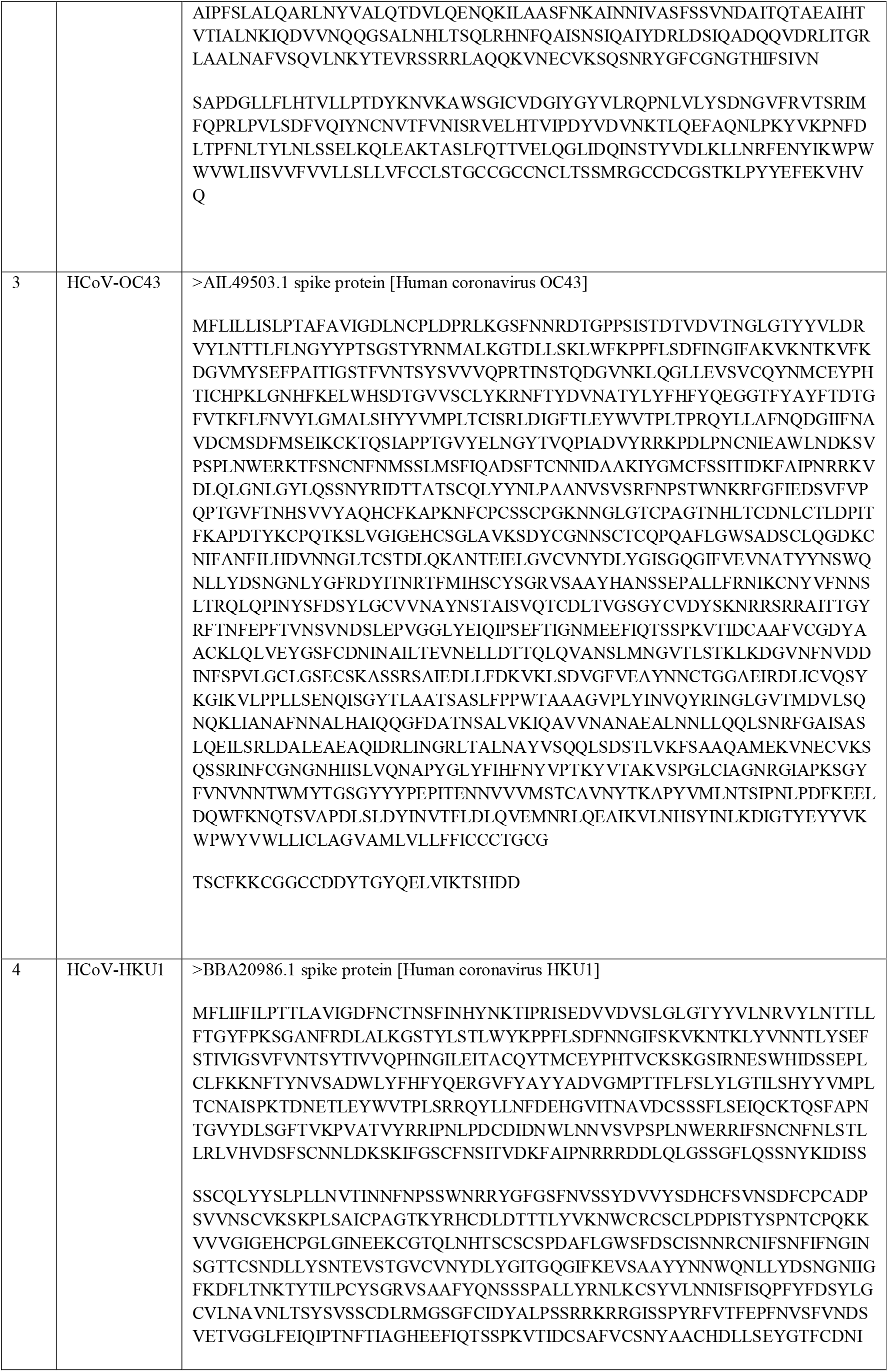
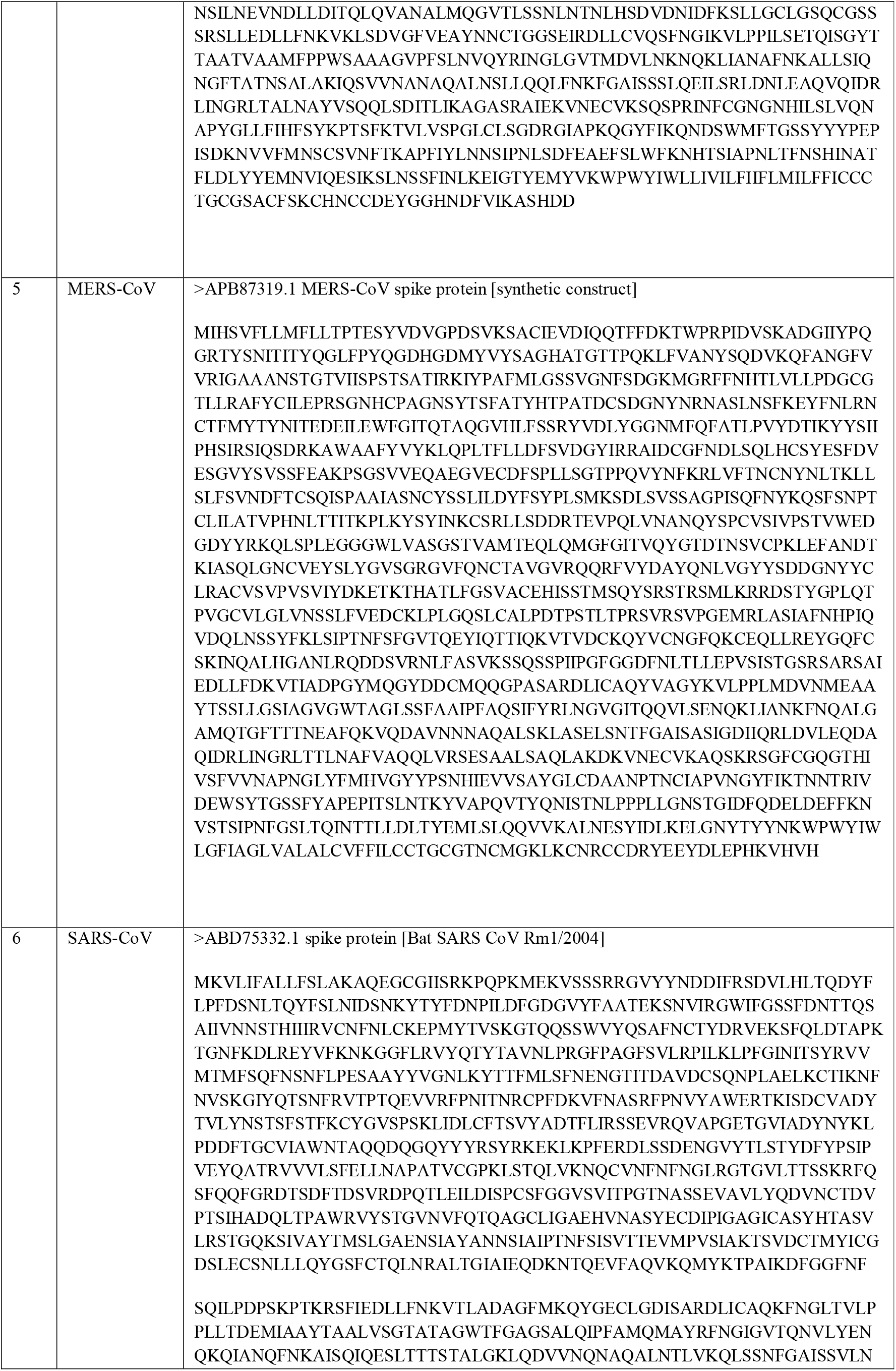
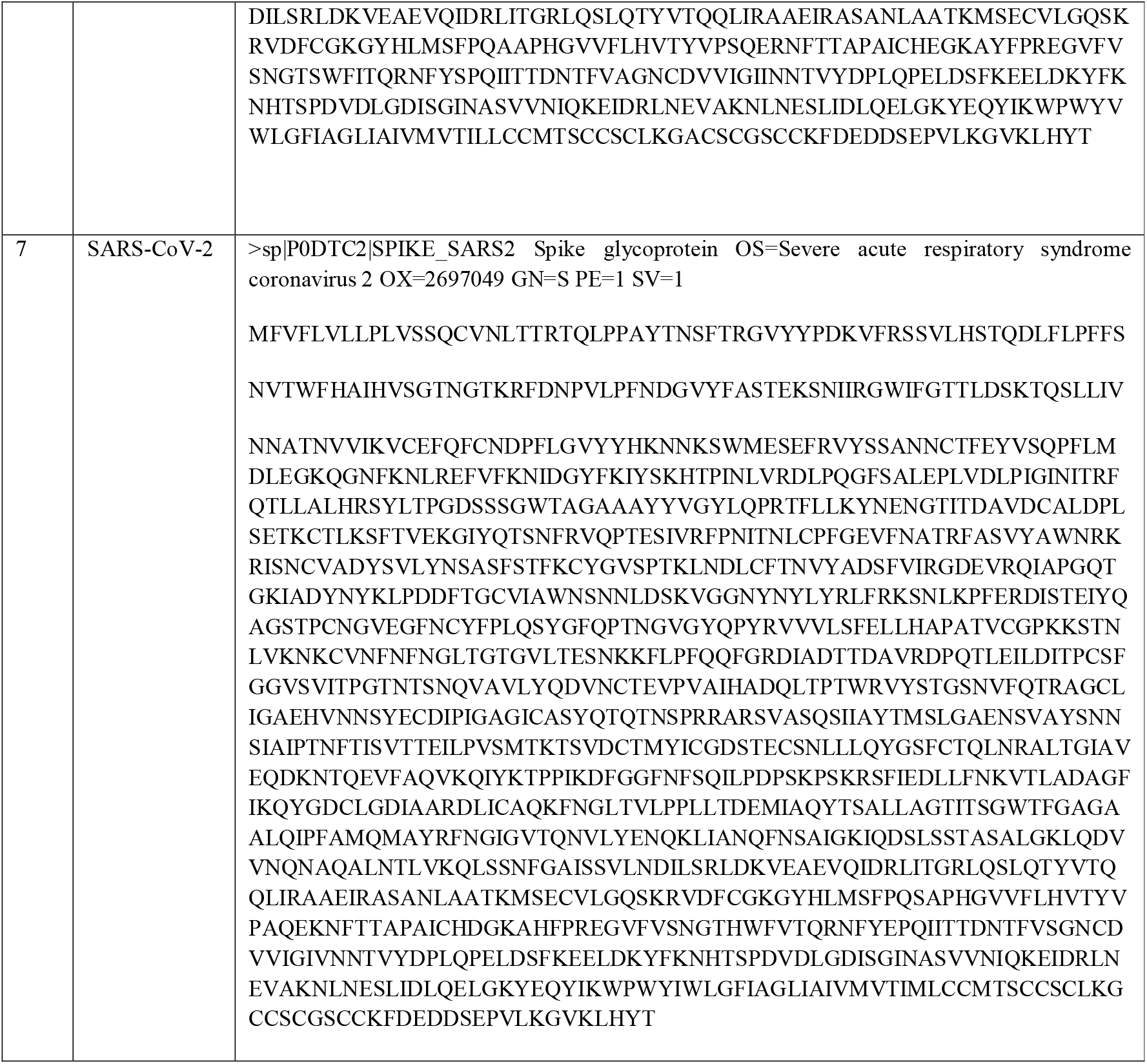
The sequences of the S protein in FASTA format as retrieved from the UniProtKB database.

### 2.3 Prediction of T-cell epitopes

Herein, we have predicted 9-mer CTL epitopes for each of the HCoV strain employing NetMHCpan4.1 web server which is also recommended by IEDB (Immune Epitope Database). The NetMHCpan-4.1 server (https://services.healthtech.dtu.dk/service.php?NetMHCpan-4.1) employs artificial neural networks (ANNs) to predict the binding of epitopes to any MHC molecule of known sequence. Its prediction method has been developed using a combined training set of >850,000 quantitative BA (Binding Affinity) and Mass-Spectrometry EL (Eluted Ligands) peptides(Reynisson, Alvarez, Paul, Peters, & Nielsen, 2020). In the present study, HLA-A*24:02 supertype of human leukocyte antigen (HLA) allele has been employed for all the predictions. The rationale behind selecting this supertype allele was the availability of high-resolution X-ray crystal structure (PDB ID 3I6L) of HLA-A*24:02 in complex with an epitope N1 derived from SARS-CoV N protein. The other reason behind the selection of this supertype was that according to Lu et al., one of the most common HLA-A allele found in the Asian population is HLA-A*24:02(Lu et al., 2005).

Best two 9-mer peptides from each HCoV strain was selected based on their binding affinity (nM) and percentile rank. Thus, a total of 14 CTL epitopes were selected for further study. But we wanted to refine our selection of CTL epitopes further and hence these epitopes were put for epitope conservancy analysis by employing the IEDB tool for the epitope conservancy analysis(Bui, Sidney, Li, Fusseder, & Sette, 2007a). For the present work linear epitope conservancy analysis was done with a sequence identity threshold of 100%. The epitopes with the highest minimum sequence identity were taken up for further analysis. Further, these 14 epitopes were put for molecular docking studies using peptide docking under the Biologics suite of Maestro (Release 2018-3). The X-ray crystal structure of HLA-A*24:02 protein (PDB ID 3I6L) was retrieved from the protein data bank (PDB)(Berman et al., 2000; Liu et al., 2010). The retrieved protein structure was prepared using the protein preparation wizard of Maestro and the optimised protein structure was energetically minimized by employing OPLS3e force field(Kumar, Rathi, & Kini, 2019; Madhavi Sastry, Adzhigirey, Day, Annabhimoju, & Sherman, 2013). The epitope sequences were imported in the workspace and using peptide builder peptides were generated. The generated peptides were energetically minimized and docked against the receptor grid generated at the binding site of epitope N1 derived from SARS-CoV N protein on the HLA-A*24:02 protein. The number of poses to return for each docking run was fixed at 10 as peptides are very flexible hence, several docking runs were used for each peptide. Each docking run employed different conformation of the epitope to enhance the sampling(Tubert-Brohman, Sherman, Repasky, & Beuming, 2013). The MM-GBSA (molecular mechanics generalized Born model and solvent accessibility) was used as the scoring method which is more accurate than the Glide score method but less fast(J. Li et al., 2011).

The epitopes were ranked based on the docking score and the highest-ranked CTL epitope was further put for molecular dynamics (MD) simulations analysis by using Desmond module of Maestro suite(Bowers et al., 2006). The docked complex of the selected epitopes with the HLA-A*24:02 protein (PDB ID 3I6L) was first put for system builder where the Na^+^ and Cl^-^ were added to neutralize the charges. SPC (simple point charge) was used as the solvent system in an orthorhombic boundary box shape. The solvated system of epitope and the HLA-A*24:02 protein was put for minimization by using SD (steepest descent) method. The convergence threshold was fixed at 1 kcal/mol/Å with a maximum iteration of 2000. Slow relaxation protocol was followed for equilibration at a temperature of 300 K and pressure of 1.01325 bar. Nose–Hoover thermostat was used to maintain the temperature while Martina–Tobias–Klein was the barostat employed(Kumar, Rai, Rathi, Agarwal, & Kini, 2020). The minimized protein-epitope complex was simulated for 50 ns to analyse the stability of the epitope-protein complex under the biological conditions. Based on the above methodologies i.e. percentile rank, binding affinity, minimum percentage conservancy and docking score, a total of ten CTL epitopes were selected for the design of vaccine candidate

### 2.4 Prediction of linear B-cell epitope and Helper T-cell (HTL) epitopes

B-cell epitopes are a specific part of the antigen recognized by the B lymphocytes of our immune system. For the present work, BepiPred-2.0 webserver (http://www.cbs.dtu.dk/services/BepiPred/) has been employed to predict linear B-cell epitopes of 14 sequence length for all the seven HCoV strains. BepiPred-2.0 is superior to other sequence-based linear epitope prediction tools and employs RF (random forest) algorithm trained on peptides annotated from antigen-antibody protein complexes(Jespersen, Peters, Nielsen, & Marcatili, 2017). HTL epitopes of 15 sequence length were predicted using the NetMHCII 2.2 Server (http://www.cbs.dtu.dk/services/NetMHCII/) for all the seven strains. The NetMHCII 2.2 server employs artificial neuron networks to predict the binding of epitopes to HLA-DQ, HLA-DP, and HLA-DR alleles(Nielsen & Lund, 2009).

### 2.5 Multi-epitope vaccine candidate construction and its conservancy analysis

The multi-epitope vaccine candidate was designed by linking different epitopes and adjuvants. CTLs epitopes were linked through an AAY linker while HTL and B-cell epitope was linked using GPGPG linker. At the N-terminal, a 50 S ribosomal protein L7/L12 (Locus RL7_MYCTU) was added as an adjuvant (Accession no. P9WHE3) through an EAAAK linker. The adjuvant sequence was retrieved from the UniProt database (http://www.uniprot.org/). At the carboxy-terminal end, a 6x-His tail was also added to generate a final vaccine construct. The final designed vaccine peptide was put for conservancy analysis without the adjuvant across the seven strains of the selected S-protein sequences of the HCoVs(Bui, Sidney, Li, Fusseder, & Sette, 2007b).

### 2.6 Prediction of IFN-gamma inducing epitope

Cytokines like Interferon-gamma (IFN-γ) play a crucial role in the immune responses by stimulating NK (natural killer) cells and macrophages and provide an enhanced response to MHC antigen. Herein we have employed the IFNepitope server (http://crdd.osdd.net/raghava/ifnepitope/scan.php) to predict 15-mer IFN-γ epitopes for the designed vaccine candidate. IFN-γ epitopes were predicted separately for the adjuvant and the main vaccine peptide due to the limitation on the number of sequences that can be used as input in the server. The server employs support vector machine (SVM) hybrid approach for the prediction of IFN-γ epitopes(Dhanda, Vir, & Raghava, 2013).

### 2.7 Antigenicity, allergenicity and assessment of physicochemical properties of the vaccine construct

Antigenicity prediction for the vaccine construct was carried out using ANTIGENpro (http://scratch.proteomics.ics.uci.edu/) and VaxiJen2.0 server (http://www.ddg-pharmfac.net/vaxijen/VaxiJen/VaxiJen.html). Like ANTIGENpro the antigenicity prediction by VaxiJen v2.0 is alignment-free and employs the physiochemical parameters of the protein(Doytchinova & Flower, 2007). Allergenicity of the designed vaccine construct was predicted using AllerCatPro (https://allercatpro.bii.a-star.edu.sg/) and AllergenFP (http://ddg-pharmfac.net/AllergenFP/) servers. AllerCatPro predicts the allergenicity of the query proteins by comparing their 3D structure along with their amino acid sequence with a dataset of 4180 unique protein sequences which have been reported to be allergenic(Maurer-Stroh et al., 2019). To predict the allergenicity of proteins, AllergenFP employs a descriptor-based fingerprint approach and thus is an alignment-free tool(Dimitrov, Naneva, Doytchinova, & Bangov, 2014). The physiochemical parameters like theoretical pI, in vitro and in vivo half-life, amino acid composition, instability index, molecular weight, aliphatic index and GRAVY (grand average of hydropathicity) of the vaccine construct was analysed by the ProtParam (http://web.expasy.org/protparam/) web server(Gasteiger et al., 2005). The solubility of the vaccine construct was analysed by the Protein-Sol (https://protein-sol.manchester.ac.uk/) web server(Hebditch, Carballo-Amador, Charonis, Curtis, & Warwicker, 2017).

### 2.8 Secondary and tertiary structure prediction of the vaccine construct

The secondary structure of the designed vaccine construct was predicted by employing PSIPRED (http://bioinf.cs.ucl.ac.uk/psipred/) and RaptorX Property (http://raptorx.uchicago.edu/StructurePropertyPred/predict/) web servers. PSIPRED 4.0 performed Position-specific iterated BLAST (Psi-Blast), for the identification of sequences which showed considerable homology to the vaccine peptide. These sequences were selected to build a position-specific scoring matrix. The RaptorX Property web server employs a template-free approach to predict the secondary structure properties of the query protein. Its algorithm is based on DeepCNF (Deep Convolutional Neural Fields) to simultaneously predict secondary structure (SS), disorder regions (DISO) and solvent accessibility (ACC)(S. Wang, Peng, Ma, & Xu, 2016). The tertiary structure of the designed multi-epitope vaccine candidate was predicted by the I-TASSER server (https://zhanglab.ccmb.med.umich.edu/I-TASSER/) webserver. The I-TASSER (Iterative Threading Assembly Refinement) server is an integrated platform to predict 3D structure form a peptide sequence. It generates 3D atomic models from multiple threading alignments and iterative structural assembly simulations and has been recognized as the best server for the prediction of protein structure(Roy, Kucukural, & Zhang, 2010). The best 3D-model for the vaccine construct obtained from I-TASSER web server was refined first by the ModRefiner (https://zhanglab.ccmb.med.umich.edu/ModRefiner/) and then by using the GalaxyRefine server (http://galaxy.seoklab.org/cgi-bin/submit.cgi?type=REFINE). The refinement of protein structures by the ModRefiner server is based on a two-step, atomic-level energy minimization which leads to the improvements in both local and global structures(Xu & Zhang, 2011). GalaxyRefine method first rebuilds side chains and then carries out side-chain repacking and finally employs MD simulation for overall structural relaxation(Heo, Park, & Seok, 2013). Finally, the refined 3D model of the vaccine construct was validated using ProSA-web (https://prosa.services.came.sbg.ac.at/prosa.php) and ERRAT server (http://services.mbi.ucla.edu/ERRAT/). Ramachandran plot was obtained from protein structure quality tool of the Prime module in Maestro11.4.

### 2.9 Protein-protein docking studies

Molecular docking was carried out to understand the binding of the vaccine candidate with TLR-3 receptor as the immune response depends upon the interaction between an antigenic molecule (here the designed vaccine candidate) and an immune receptor (in this case TLR-3). The first step in this milieu was the prediction of the binding sites of the proteins which were carried out using the CASTp (http://sts.bioe.uic.edu/castp/) server(Binkowski, Naghibzadeh, & Liang, 2003). Molecular docking of the vaccine candidate with the TLR3 (PDB ID: 1ZIW) receptor was carried out using the HADDOCK 2.4 web server(Van Zundert et al., 2016) (https://wenmr.science.uu.nl/haddock2.4/) and GRAMM-X Simulation web server(Tovchigrechko & Vakser, 2006) (http://vakser.compbio.ku.edu/resources/gramm/grammx/). The interactions were visualized using LIGPLOT v.4.5.3 (https://www.ebi.ac.uk/thornton-srv/software/LIGPLOT/)(Wallace, Laskowski, & Thornton, 1995). Finally, the binding energy of the top-ranked docking pose of the vaccine candidate-TLR4 complex was predicted using the PRODIGY webserver https://nestor.science.uu.nl/prodigy/)(Xue, Rodrigues, Kastritis, Bonvin, & Vangone, 2016).

### 2.10 Prediction of discontinuous B-cell epitopes

It has been reported that more than 90% of B-cell epitopes are discontinuous, i.e. they comprise of distantly separated segments in the pathogen protein sequence which are brought close by the folding of the protein(Barlow, Edwards, & Thornton, 1986). Herein, we employed ElliPro (http://tools.iedb.org/ellipro/) for the prediction of discontinuous B-cell epitopes of the validated 3D structure of the vaccine candidate(Ponomarenko et al., 2008).

### 2.11 *In silico* cloning and codon optimization of the vaccine construct

Java Codon Adaptation Tool (JCat) server (http://www.prodoric.de/JCat) was employed for reverse translation and codon optimization so that the designed multi-epitope vaccine candidate could be expressed in a selected expression vector(Grote et al., 2005). Codon optimization was carried out to express the final vaccine candidate in the *E. coli* (strain K12) host. The output of the JCat server comprises of the codon adaptation index (CAI) and percentage GC content. CAI score reflects codon usage biases and ideally, the value of CAI should be 1.0 but a score above 0.8 is a good score(Grote et al., 2005). The gene sequences of the designed vaccine candidate were optimized and two restriction sites (*Nde* I and *Xho*) were introduced at the C and N-terminals of the sequence. In the last step, the optimized gene sequence of the vaccine construct along with the inserted restriction sites was inserted into the pET-28a (+) vector using SnapGene programme.

### 2.12 Immune simulation

*In silico* immune simulations were carried out for further characterization of the immunogenicity and immune response profile of the designed multi-epitope vaccine candidate. Herein, we employed the C-ImmSim server (http://150.146.2.1/C-IMMSIM/index.php) for immune simulations studies which is an agent-based model that employs a position-specific scoring matrix (PSSM)(Rapin, Lund, Bernaschi, & Castiglione, 2010). Three injections were given at four weeks intervals by keeping all simulation parameters at default setting with time steps set at 1, 84, and 168 (each time step is 8 hours and time step 1 is injection at time = 0).

## 3. Results and Discussion

### 3.1 Prediction of T-cell epitopes

The goal of T-cell epitope prediction is to identify peptides with shortest sequence length within an antigen which is capable of stimulating either CD4 or CD8 T-cells(Ahmed & Maeurer, 2009). Antigen-presenting cells (APCs) present T-cell epitopes on the surface where they bind to either MHC class I protein or MHC-II proteins. It has been reported that MHC-I protein present T-cell epitopes of 8-11 amino acid residue length while MHC-II presents longer peptides in the range of 13-17 amino acid residues(Steers et al., 2014). T-cell epitopes bound to MHC I protein are recognized by CD8 T-cells which become CTL (cytotoxic T lymphocytes), while those presented by MHC-II are recognized by CD4 T-cells and become helper T-cells(Sanchez-Trincado, Gomez-Perosanz, & Reche, 2017). Many authors have reported these tools for CTL epitope prediction not only against SARS-CoV-2 but also against SARS-CoV(Abraham Peele et al., 2020; Enayatkhani et al., 2020; Naz, Shahid, Butt, Awan, & Ali, 2020; Oany et al., 2014; Panda et al., 2020). But this is the first such analysis involving seven strains of the HCoVs. Our present hypothesis to screen the S protein sequences of all the seven reported HCoVs for identifying CTL epitopes as a potential vaccine candidate against SARS-CoV-2 is also supported by the recent findings of Barun et al., where they have demonstrated the presence of S-cross-reactive T cells in unexposed healthy individuals probably because of previous exposure to endemic coronaviruses like 229E or OC43(Braun et al., 2020).

A total of 142 CTL (9-mer) epitopes were predicted to be strong binders for the seven selected S protein of different HCoVs using NetMHCpan4.1 server with default settings. We selected the best two epitopes based on their percentile rank, binding affinity (nM) and binding level for each of the HCoV strain. Thus, a total of fourteen 9-mer CTL epitopes were selected for further study. The conservancy analysis of these fourteen CTL epitopes showed that QYIKWPWYI epitope from SARS-CoV-2 strain had a minimum conservancy of 66.67% across all the strains. Another epitope with sequence YYNKWPWYI from MERS strain showed a minimum conservancy of 55.56% across all the strains. Hence, these two CTL epitopes were selected for the final vaccine design. Peptide docking analysis of these epitopes with HLA-A*24:02 protein showed that these peptides showed favourable binding with the protein. But we selected only one CTL epitope (NYNYLYRLF) which showed the highest docking score of −12.759. MD simulations study for this epitope showed that the epitope-protein complex was stable throughout the simulations period as reflected by their RMSD (root mean square deviation) plot in Fig. 2a. A histogram representing the interaction of the epitopes with the amino acid residues of the protein was also plotted as shown in Fig. 2b. The non-bonding interactions were similar to the ones observed during peptide docking studies, but water bridges were formed with LYS66, HIS70, ASP74, TYR84, THR143 and GLN155 residues. A 2D interaction diagram (Fig. 2c) suggested that apart from intermolecular H-bond interactions, there were quite several intramolecular H-bond interactions which could make the epitope-protein complex more stable. The importance of intramolecular H-bonding for the epitopes binding to HLA-A*24:02 protein has also been highlighted by Liu et al(Liu et al., 2010). Several other authors like Peele et al. and Enayatkhani et al. have also employed MD simulations studies to analyse the stability of the designed epitopes and protein complex(Abraham Peele et al., 2020; Enayatkhani et al., 2020). Hence, for the final vaccine design, a total of ten CTL epitopes (Table 2) were selected out of which seven were the best-ranked epitope from each strain, two were the best-conserved epitope across all the strains and one was the best-docked epitope with HLA-A*24:02 protein.

**Table 2.**
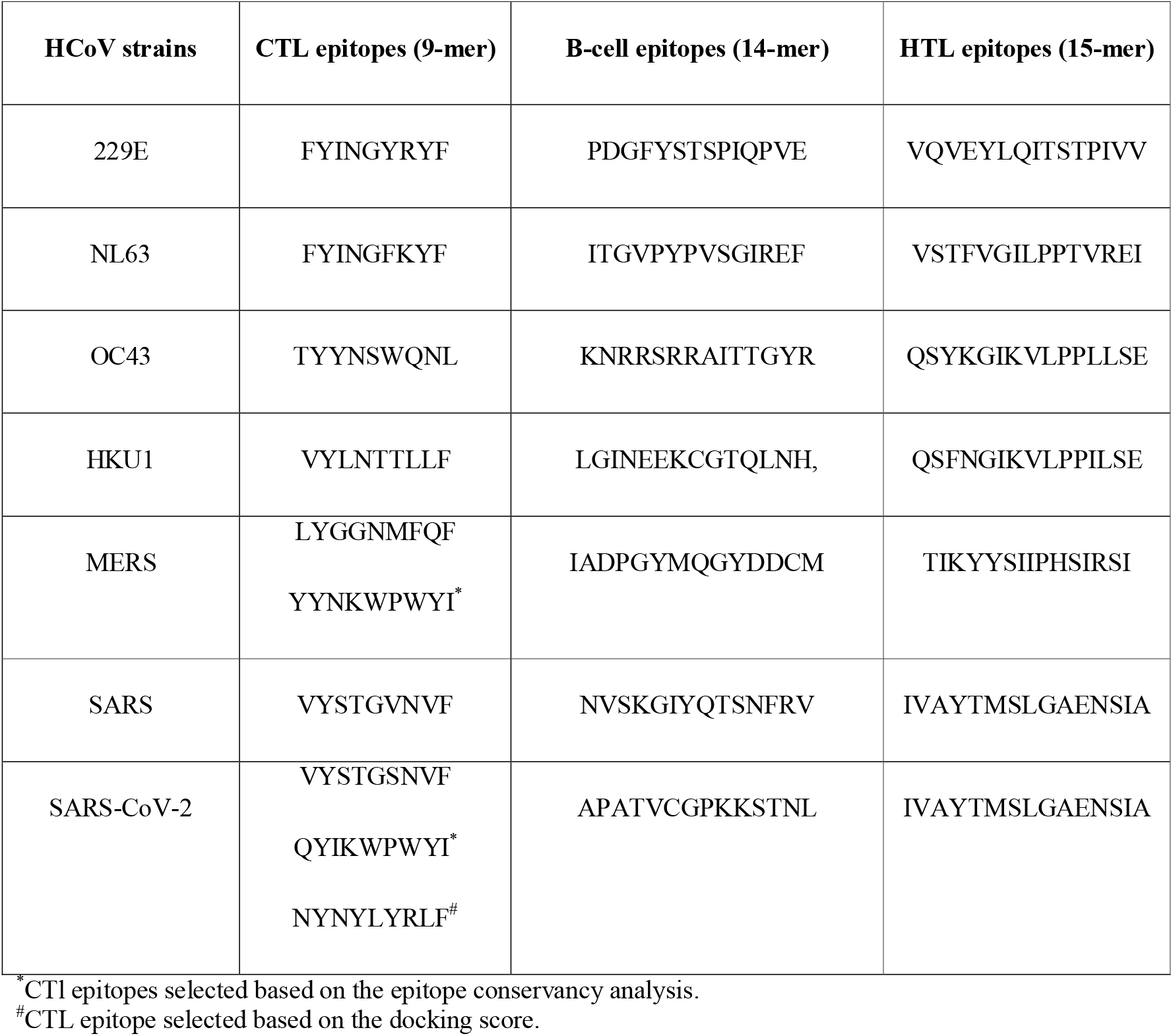
List of the selected CTL, HTL and linear B-cell epitopes from S protein sequences of different HCoV strains for constructing the multi-epitope vaccine candidate.

**Fig. 2.**
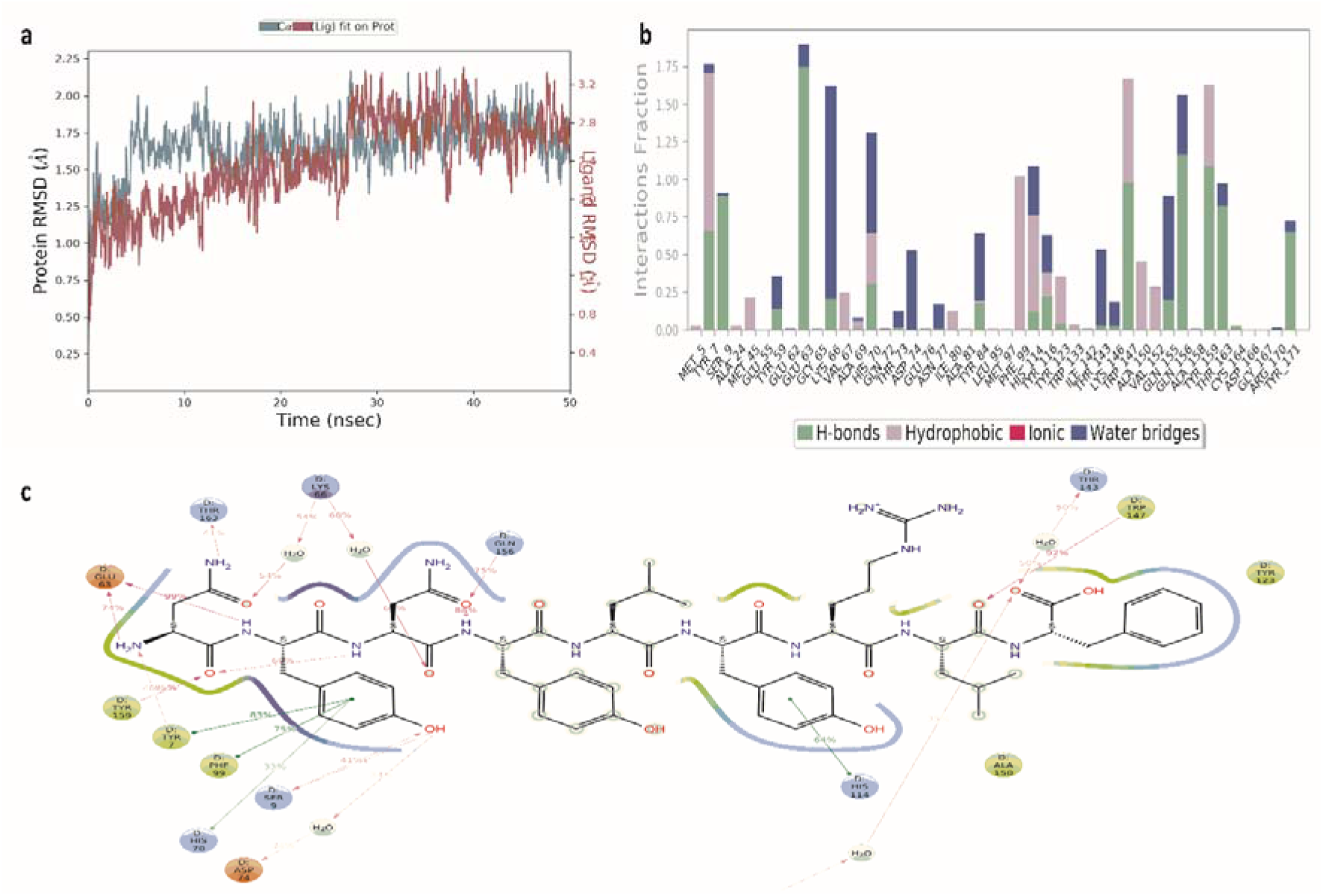
MD simulations report of the docked complex of CTL-epitope “NYNYLYRLF” with HLA-A*24:02 protein (PDB ID 3I6L). (a) RMSD plot of CTL-epitope 2 in complex with HLA-A*24:02 protein. (b) Histogram of the CTL-epitope contact with HLA-A*24:02 protein. (c) CTL-epitope contact summary with HLA-A*24:02 protein (PDB ID 3I6L) in 2D.

### 3.2 Prediction of linear B-cell epitope and Helper T-cell (HTL) epitopes

A total of seven 14-mer B-cell epitopes (Table 2) were selected based on the epitope threshold and relative surface accessibility as predicted by BepiPred2.0 server. A total of seven 15-mer high-binding MHC-II epitopes (Table 2) for human alleles HLA-DR, HLA-DQ and HLA-DP as predicted by the NetMHCII 2.2 web server was selected. Some of the B-cell epitopes and HTL epitopes overlapped.

### 3.3 Multi-epitope vaccine candidate construction and its conservancy analysis

The final multi-epitope vaccine construct comprised of 10 CTL epitopes, 7 linear B-cell epitopes, and 7 HTL epitopes. The predicted CTL epitopes were joined by using AAY linker while B-cell epitope and TL epitopes were fused with GPGPG linkers. The 50S ribosomal L7/L12 (Locus RL7_MYCTU) was selected as the adjuvant to accentuate antigen-specific immune responses and fused at the N-terminal by using EAAAK linker. Finally, to help in protein identification and purification a 6xHis tail was inserted at the carboxy-terminal end of the vaccine peptide. Thus, the final multi-epitope vaccine candidate was constructed with a total of 532 amino acid residues derived from 24 merged multi-epitopes. The final designed vaccine peptide was put for conservancy analysis without the adjuvant across the seven strains of the selected S-protein sequences of the HCoV strains. The sequences of the vaccine candidate showed a minimum and maximum conservancy of 73.97% across all the strains of the HCoVs. A higher degree of conservation across all the HCoV strains might confer broader protection across multiple strains by recognition of the conserved epitopes by our immune system. Hence, we hypothesize that this vaccine construct which has more than 73% of sequences conserved across the seven HCoV strains might be a potential broad-spectrum vaccine candidate against HCoVs.

### 3.4 Prediction of IFN-gamma inducing epitope

Due to the restriction on the number of residues that can be used as input in the IFNepitope server, IFN-γ inducing epitopes were predicted separately for the adjuvant and main vaccine peptide. For the adjuvant, a total of 127 potential IFN-γ inducing epitopes (15-mer) were predicted. For the main vaccine peptide, 389 potential 15-mer epitopes were predicted out of which 82 had a positive prediction score.

### 3.5 Antigenicity, allergenicity and assessment of physicochemical properties of the vaccine construct

As per the predictions of the VaxiJen 2.0 server, the antigenicity of the vaccine construct along with the adjuvant sequence was found to be 0.7106 in a bacteria model at a threshold of 0.5 while the main vaccine sequence without the adjuvant showed a score of 0.8135. As per the predictions of ANTIGENpro server the probable antigenicity of the vaccine candidate with the adjuvant was 0.829721 while without adjuvant was 0.869507. This result indicates that the designed vaccine candidates are antigenic (with or without adjuvant) in nature. The main vaccine sequence seems to be more antigenic without the adjuvants as per the predicted scores. The allergenicity of the vaccine candidate was predicted by the AllerCatPro and AllergenFP servers and both predicted it to be non-allergenic. The molecular weight (MW) of the final vaccine candidate was predicted to be 56.5 kDa. The theoretical isoelectric point (pI) value was predicted to be 7.11. The half-life was predicted to be 30 hours in mammalian reticulocytes (*in vitro*), and>10 hours in E. coli and >20 hours in yeast (*in vivo*). The instability index (II) was computed to be 22.81 which classifies the vaccine candidate to be stable. It had an aliphatic index of 79.29 which suggests it to be thermostable(Ikai, 1980). The GRAVY score for the vaccine candidate was computed to be −0.407 which indicates that it could be hydrophilic in nature.(Ali et al., 2017) The predicted scaled solubility value was computed to be 0.432 which indicates that the vaccine construct would have an acceptable solubility profile.

### 3.6 Secondary and tertiary structure prediction of the vaccine construct

The secondary structure of the final multi-epitope vaccine peptide was computed, and it was predicted to have 20% alpha-helix, 20% beta-sheet and 59% coil (Fig. 3). The solvent accessibility of amino acid residues as predicted by RaptorX property server showed that 49% of the total residues were exposed, 24% were medium exposed and 26% was buried. A total of 42 residues i.e. 7% of residues were computed to be in the disordered region. Five tertiary structure models of the final vaccine construct were predicted by the I-TASSER server based on 10 threading templates, of which PDB hits 1dd4A, 5tsjN, 1rquA, 2nbiA, 1dd3A, 2ocwA and 2ftc were best hits. All of the 10 selected templates showed good alignment which was evident from their *Z-score* values which ranged 1.02 to 5.35. The first model (Fig. 4a) was selected for further refining which had an estimated C-score of 0.77, TM score of 0.62±0.14 and RMSD value 9.2±4.6Å. The *C*-score range is between −5 and 2, where higher values towards 2 indicate higher confidence. The TM-score is an indication of the structural similarity between two structures and unlike RMSD, it is not sensitive to local error(Y. Zhang & Skolnick, 2004). From the TM value of the selected model, we can infer that the model has correct topology. This selected model was further put for refining first by ModRefiner and then by GalaxyRefine server which generated five models. According to the model quality scores (Table 3) for all the five refined models, model 3 (Fig. 4b) was found to be the best model. It had a GDT-HA score of 0.9408, RMSD value 0.443, MolProbity score of 2.072, clash score of 11.3 and the poor rotamers score was 0.2. The Ramachandran plot score for model 3 was predicted to be 91.5% which was less when it was put for validation by the Ramachandran plot analysis. The Ramachandran plot of the selected model 3 (Fig. 4c) suggests that it’s 94.528% of the residues are in the favourable region with 2.64% of residues in the allowed region. The overall quality factor was computed by the ERRAT server and for the selected model 3, it was predicted to be 86.235. The ProSA-web computed the Z-score for model 3 and it was found to be −2.41 (Fig. 4d).

**Table 3.** Detailed result of the five models refined by the GalaxyRefine server.

| Model | GDT-HA | RMSD | MolProbity | Clash score | Poor rotamers | Rama favoured |
| --- | --- | --- | --- | --- | --- | --- |
| Initial | 1.0000 | 0.000 | 3.083 | 61.4 | 2.0 | 88.1 |
| MODEL 1 | 0.9314 | 0.467 | 2.126 | 12.6 | 0.5 | 91.1 |
| MODEL 2 | 0.9305 | 0.476 | 2.111 | 11.6 | 0.5 | 90.6 |
| MODEL 3 | 0.9408 | 0.443 | 2.072 | 11.3 | 0.2 | 91.5 |
| MODEL 4 | 0.9375 | 0.465 | 2.180 | 14.2 | 0.5 | 90.9 |
| MODEL 5 | 0.9337 | 0.457 | 2.122 | 12.1 | 0.5 | 90.8 |

**Fig. 3.**
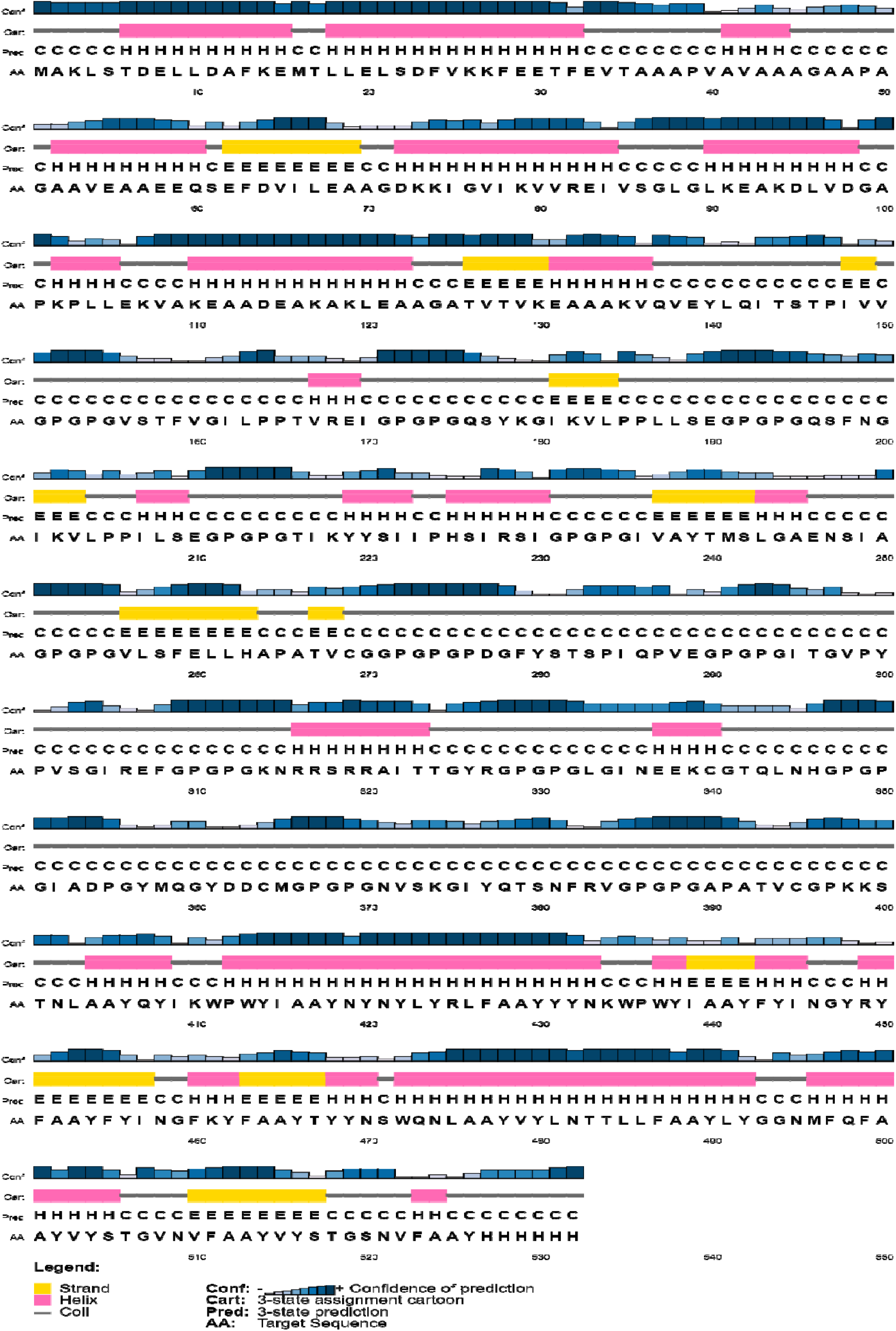
Cartoon representation of the secondary structure of the vaccine sequence as predicted by the PSIPRED.

**Fig.4.**
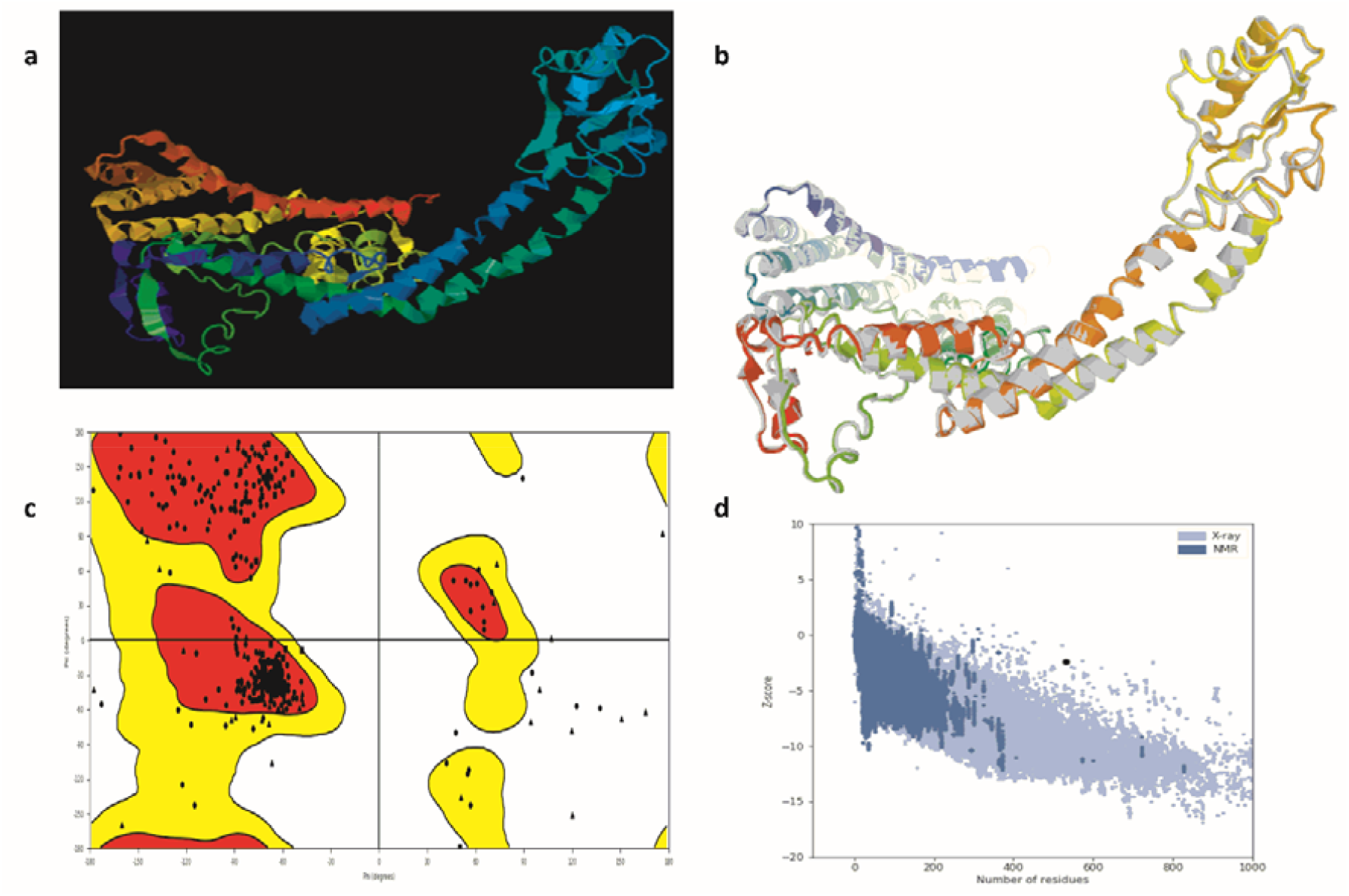
Tertiary structure modelling, refining and validation. (a) The final homology 3D model of the vaccine construct as computed by the I-TASSER. (b) Refinement of the selected I-TASSER 3D model by GalaxyRefine where the refined 3D model (coloured) has been superimposed on the initial crude model (grey). (c) Ramachandran plot of the refined 3D model of the vaccine candidate (d) Plot by ProSA-web showing the Z-score.

### 3.7 Protein-protein docking studies

The CASTp 3.0 server was employed to compute the protein binding and hydrophobic interaction sites on the surface of the TLR-3 receptor and the final refined 3D model of the vaccine construct. A total of 105 binding pockets were predicted for the vaccine candidate with different molecular surface area and volume. Similarly, for the TLR-3 protein (PDB ID 1ZIW), a total of 77 binding pockets were predicted. For the present work, a pocket (Fig. 5a) with a molecular surface area of 2125.000 Å^2^ and volume of 13438.169 Å^3^ was selected for the TLR-3 receptor. For the vaccine candidate, the selected pocket (Fig. 5b) had a molecular surface area of 917.628 Å^2^ and volume of 2414.204 Å^3^. Both the proteins were put for molecular docking simulations first using the GRAMM-X server which led to the generation of ten binding poses. The poses were visualised using the DIMPLOT of LIGPLUS version 4.5.3 and the best-docked pose of the vaccine-TLR-3 complex in 3D has been depicted in Fig. 6a. The non-bonding interactions between the docked protein complex were also visualized in 2D and have been shown in Fig. 6b. It was observed that ARG306, TYR526, ANS521 and GLY46 residues of the vaccine construct showed hydrogen bonding with TRP273, ANS247, LEU243 and HIS432 residues of the TLR-3 protein respectively. The 2D interaction plot also showed possible salt bridge between GLU307 and GLY46 of the vaccine construct with the LYS272 and SER481 of the TLR-3 protein. Further, the best-docked pose was put for the prediction of binding affinity (ΔG) and dissociation constant (K_d_). The binding energy (ΔG) for the vaccine-TLR-3 complex was computed to be −20.8 kcal/mol with a K_d_ value of 5.9 × 10^16^ M at 25.0 □. The Prodigy server also calculated the number of interfacial contacts (ICs) per property within the threshold distance of 5.5 □ and its ICs charged-charged, ICs polar-polar, and ICs apolar-apolar was computed to be 14, 7 and 64 respectively. It suggests that most of the interfacial contacts were hydrophobic in nature.

**Fig. 5.**
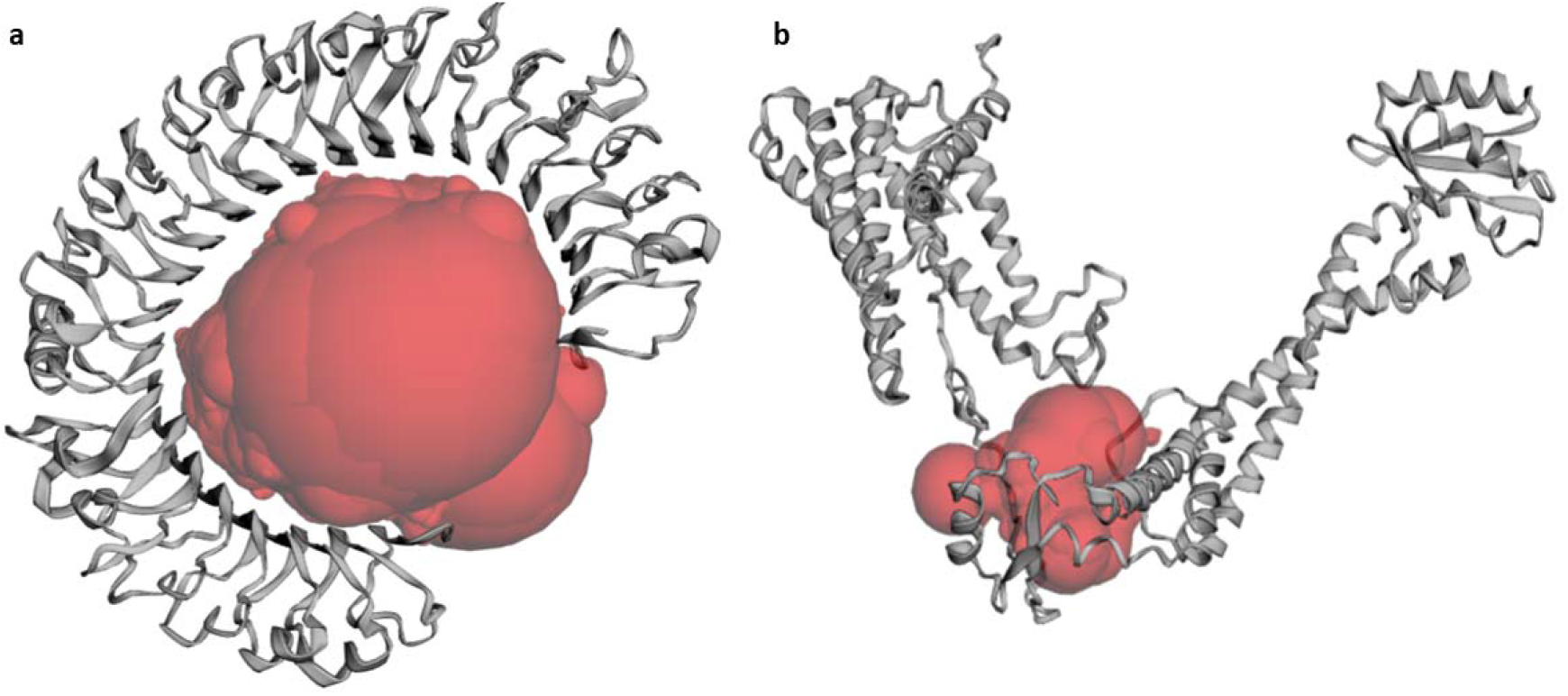
The selected binding pocket of the proteins as predicted by the CASTp3.0 server. (a) The predicted binding pocket of the TLR-3 protein. (b) The predicted binding pocket of the vaccine candidate.

**Fig. 6.**
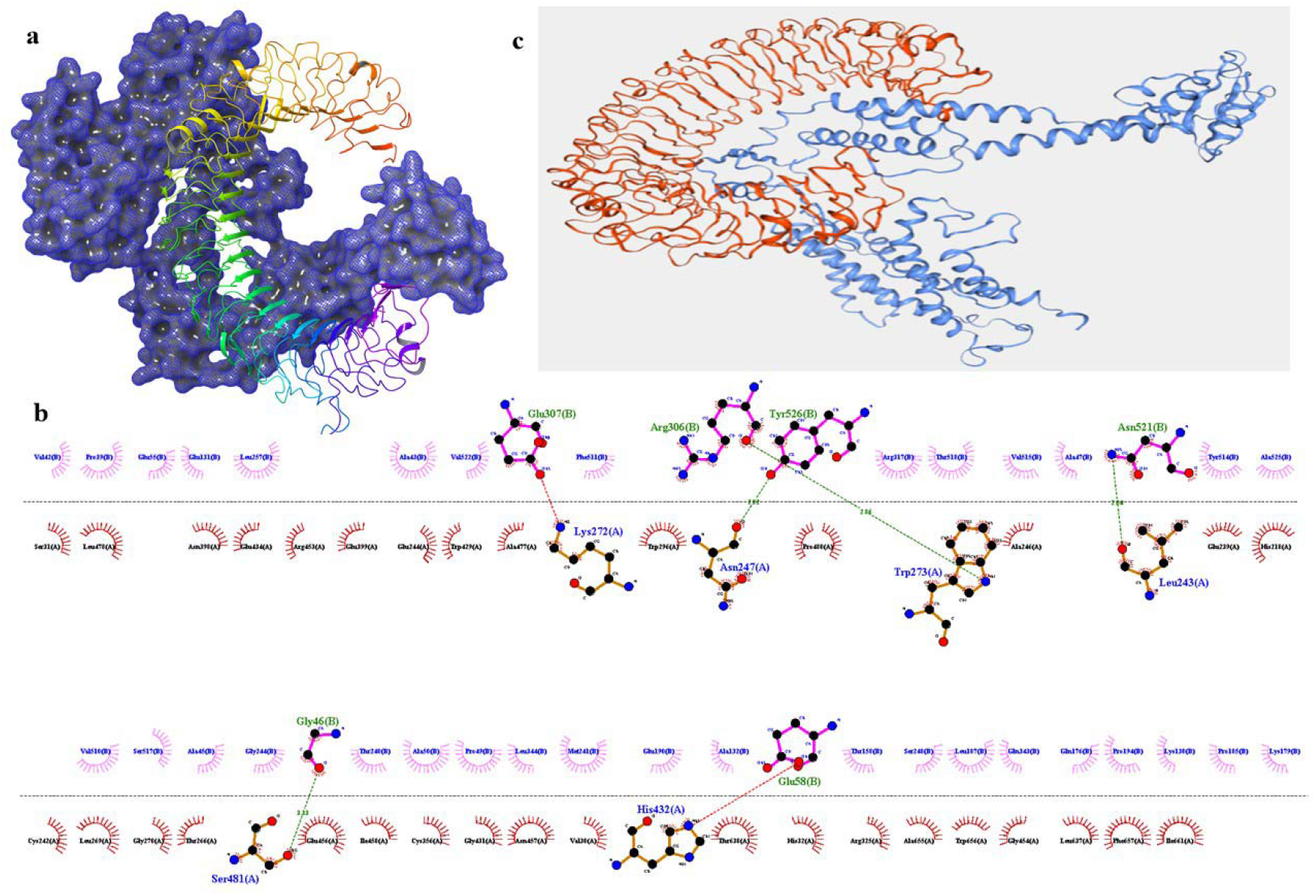
Visualization of the best-docked poses for the vaccine-TLR-3 receptor complex. (a) 3D-interaction diagram of the protein-protein complex generated by the GRAMM-X server where blue mesh structure represents the vaccine candidate while coloured ribbon shows the TLR-3 receptor. (b) 2D-interaction diagram of the protein-protein complex generated by the GRAMM-X server where chain A is from the TLR-3 receptor and chain B is the vaccine construct. (c) 3D-interaction diagram of the protein-protein complex generated by the HADDOCK2.4 server where blue ribbon represents the vaccine candidate while orange ribbon shows the TLR-3 receptor.

The HADDOCK2.4 server clustered 103 structures in 14 clusters, which represent 51% of the water-refined model generated by the HADDOCK. The clusters were ranked on several parameters and out of the 14 clusters, cluster 3 (cluster size 10) was reported to be the best cluster. It had a HADDOCK score of −79.4 +/-7.1, RMSD from the overall lowest-energy structure was 43.8 +/-0.4, Van der Waals energy −100.6 +/-15.7 kcal/mol, Electrostatic energy −415.7 +/-78.8 kcal/mol, Desolvation energy −27.2 +/-4.0 kcal/mol and Z-Score −1.5. The detailed result for all the clusters has been depicted in Fig. 7 and Fig. 8. The best pose from cluster 3 was also put for the prediction of binding affinity (ΔG) and dissociation constant (K_d_). The binding energy (ΔG) for was computed to be −20.3 kcal/mol with a K_d_ value of 1.3 × 10^15^M at 25.0 □ which was similar to that of the best docked pose generated by the GRAMM-X server. The ICs charged-charged, ICs polar-polar, and ICs apolar-apolar were computed to be 16, 7 and 25 respectively.

**Fig. 7.**
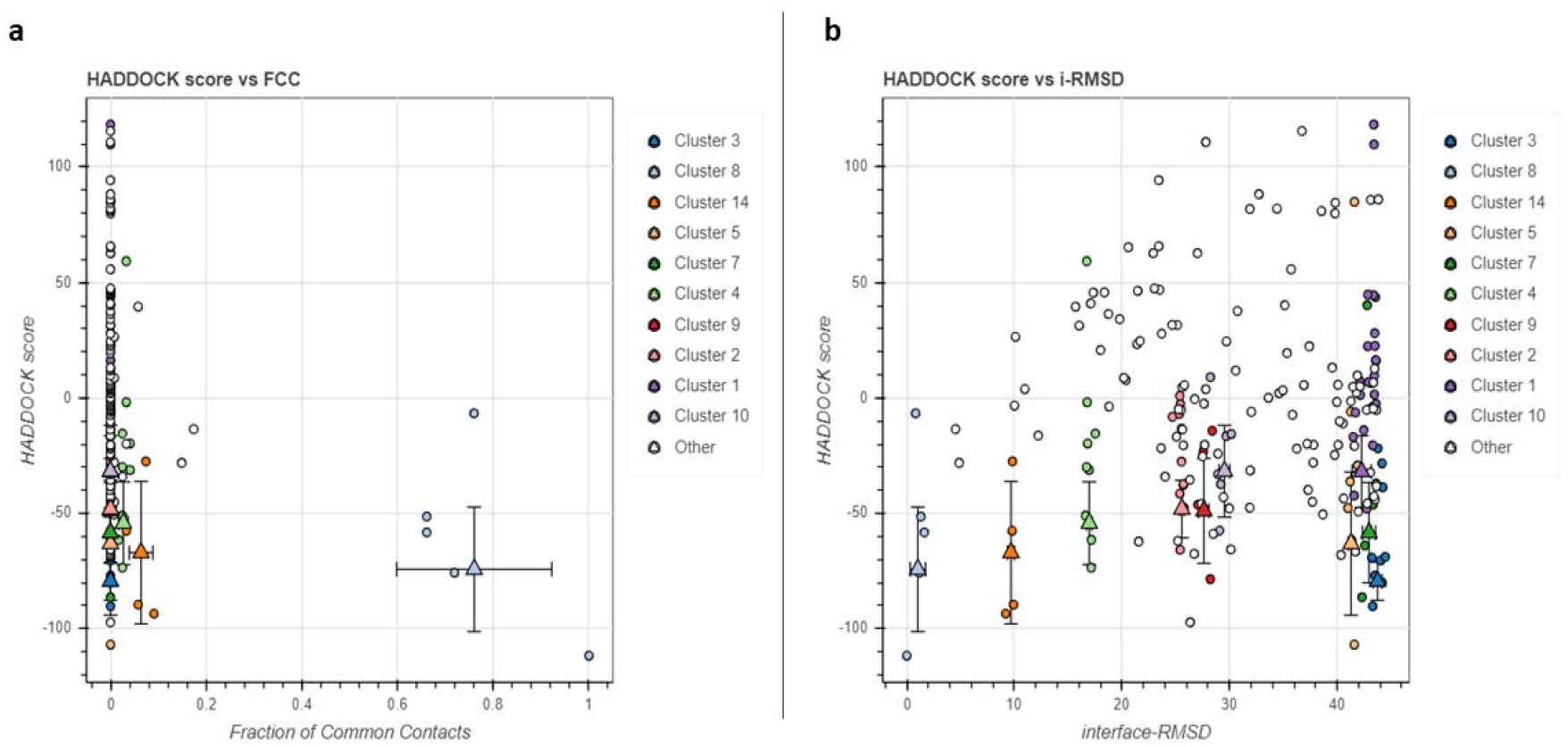
Bokeh plot showing the result of various parameters for all the clusters by the HADDCOK2.4 server. (a) Plot between HADDOCK score and FCC (fraction of common contacts). (b) Plot depicting HADDCOK score vs interface-RMSD.

**Fig. 8.**
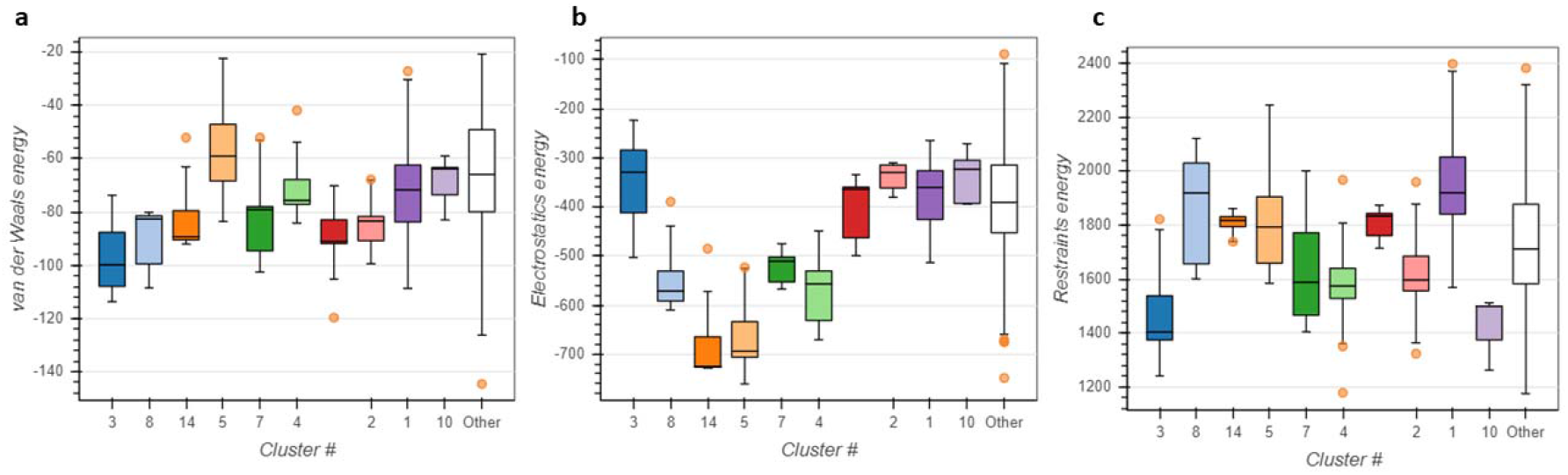
Bokeh plot depicting the analysis of various energy parameters for all the clusters as predicted by the HADDOCK2.4 server. (a) Van der Waals energy (b) Electrostatic energy (c) Restraints energy.

### 3.8 Prediction of discontinuous B-cell epitopes

The ElliPro server predicted a total of 7 discontinuous B-cell epitopes of the validated 3D structure of the vaccine candidate comprising of 270 residues. The conformational B-cell epitopes had scores ranging from 0.598 to .875 where the best epitope had only three residues while the last ranked epitope comprised of 8 residues. This study suggests that the designed multi-epitope vaccine might lead to heightened immune response though the conformational B-cell epitopes also.

### 3.9 *In silico* cloning and codon optimization of the vaccine construct

Immunoreactivity through serological assays are needed to validate a designed vaccine candidate and the first step for it is the expression of the multi-epitope vaccine candidate in a suitable host(Gori, Longhi, Peri, & Colombo, 2013). *Escherichia coli* expression systems have been widely used to produce recombinant proteins and hence to achieve maximal protein expression, we selected *E. coli* (strain K12) system for the codon optimization by the Java Codon Adaptation Tool (JCat)(Chen, 2012; Rosano & Ceccarelli, 2014). The optimized codon sequence for the final vaccine candidate was 1596 with a CAI value of 0.9237 and 53.57% of GC content. Ideally, the GC content should be in the range of 30% to 70% and hence, our vaccine candidate might show high-level expression in the bacterial host system. Finally, the optimized gene sequences of the designed vaccine candidate with two restriction sites (*Nde* I and *Xho*) at the C and N-terminals of the sequence were inserted into the pET-28a (+) vector using SnapGene software to generate the sequences of the recombinant plasmid (6892 bp) as shown in Fig. 9.

**Fig. 9.**
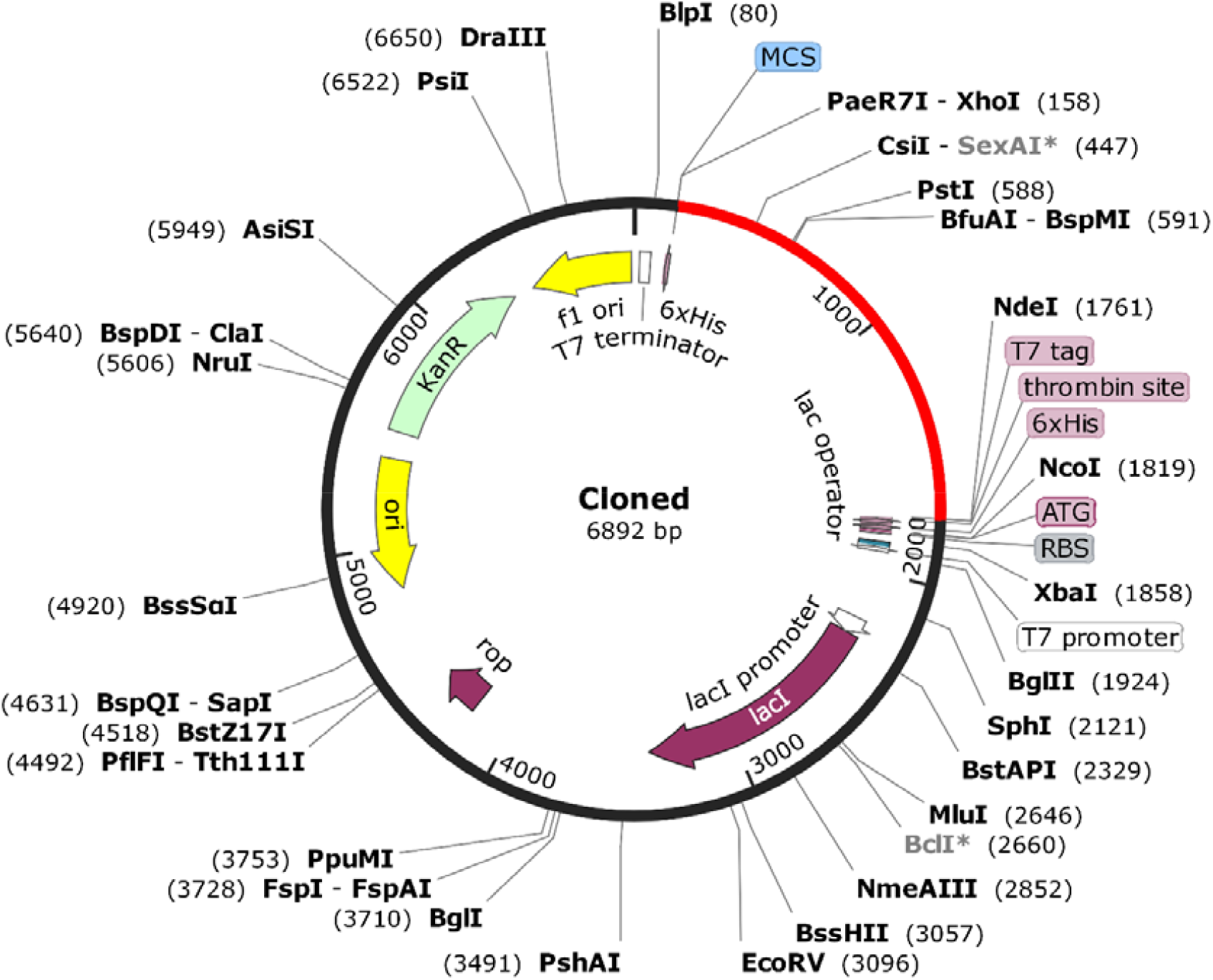
*In silico* cloning of the final multi-epitope vaccine candidate into the pET28a (+) expression vector where the red portion depicts the gene coding region for the vaccine and the black circle shows the vector backbone.

### 3.10 Immune simulation

The immune simulation by the C-ImmSim server showed results which are consistent with actual immune responses. The primary response after the first dose of the vaccine was characterized by the increased levels of IgM as shown in Fig. 10a. After the second and third dose of vaccine, there was a remarkable increase in the levels of IgG1, IgG1 + IgG2, IgM, and IgG + IgM antibodies (Fig. 10a). The level of IgM peaked after the third dose to around 2,30,000 which suggests a strong humoral immune response which is better than reported by Peele et al. for multi-epitope vaccine design against SARS-CoV-2(Abraham Peele et al., 2020) and Shey et al for multi-epitope vaccine candidate against onchocerciasis(Shey et al., 2019). It was also observed that after the second exposure the population of the antigen was also falling which indicates increased clearance by the immune system. B-cell population (Fig. 10b) and especially the B-memory cell population (>800 cells per mm^3^) also increased and peaked after the third dose. Similarly, a greater response was observed in the cytotoxic T-cells (Fig. 10c) and helper-T cell populations (Fig. 10d) with corresponding memory development which lasted for many months. It was also observed that the levels of IFN-γ and IL-2 (Fig. 10e) increased significantly after the first dose of the vaccine and maintained their peak levels after multiple exposures to the antigen. The Simpson index, D which indicates a measure of diversity, predicted this vaccine candidate to have a diverse immune response. This diverse immune response might be possible as the vaccine comprises of different epitopes.

**Fig. 10.**
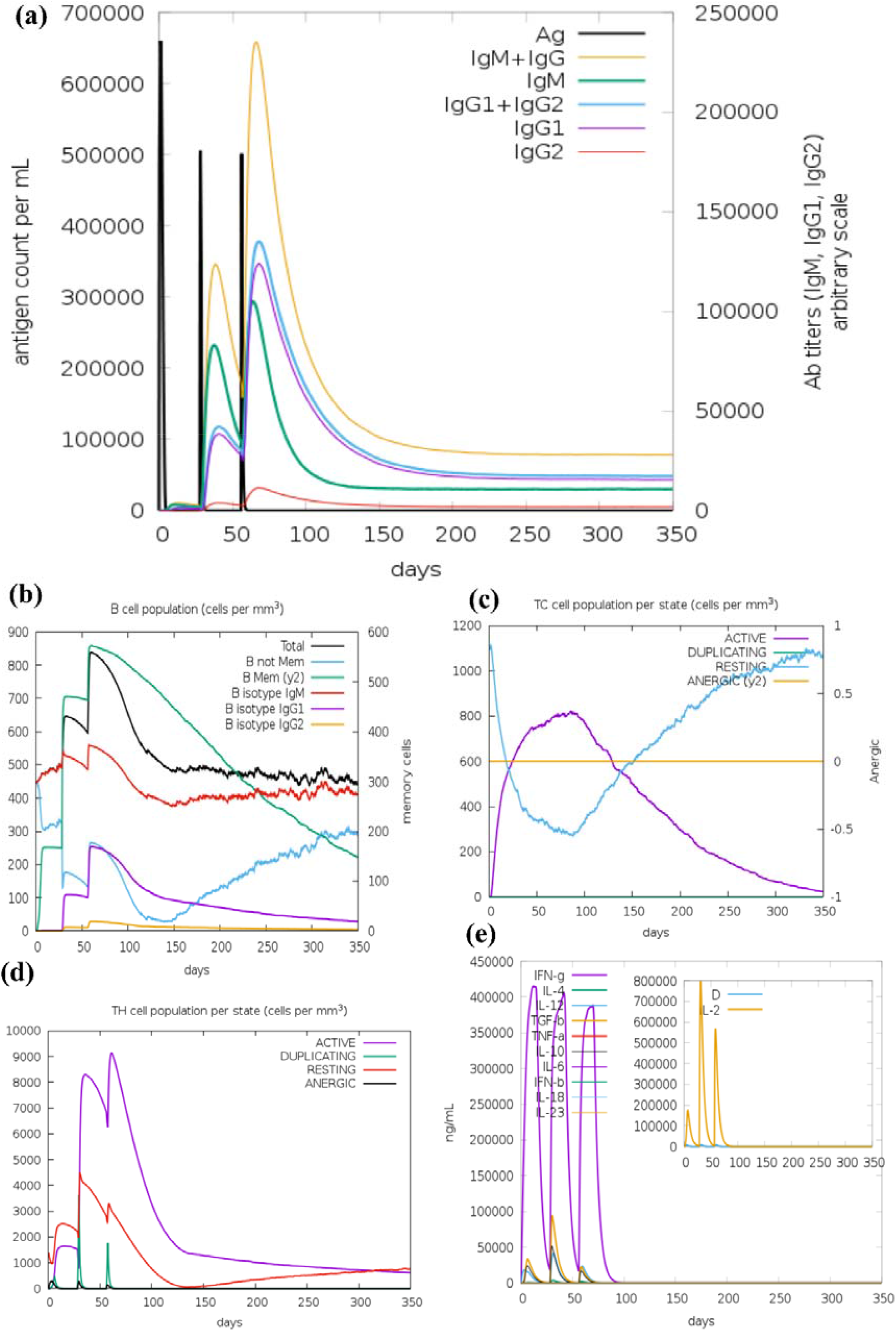
Results of immune simulation by the C-ImmSim server. (a) Immunoglobulin and immunocomplexes production in response to antigen injections at different time intervals (black vertical lines); specific subclasses have been shown as coloured peaks. (b) The increase in the populations of B-cell post three doses of the vaccine. (c) The evolution of T-cytotoxic cell population per state post three vaccine doses. (d) The evolution of T-helper cell population per state post three vaccine doses. (e) The main plot depicts the levels of cytokine levels post three vaccine doses. The insert plot depicts IL-2 level with the Simpson index, D which is indicated by the green dotted line.

## 4. Conclusions

In the present study, several machine learning based in silico tools were used to design a potential broad-spectrum multi-epitope vaccine candidate against spike protein of human coronaviruses. To the best of our knowledge, it is one of the first study, where multiple B-cell epitopes and T-cell epitopes (CTL and HTL) were predicted from the spike protein sequences of all the seven known human coronaviruses. The final vaccine candidate was found to have a minimum conservancy of 73.97% across all the seven HCoV strains and it might provide protection from SARS-CoV-2 as well as other HCoVs. The designed vaccine is predicted to be antigenic and non-allergenic. Its physicochemical properties are also in the acceptable range. Molecular docking of the refined tertiary structure of the vaccine candidate with the TLR-3 protein also indicated its favourable binding with the TLR-3 receptor. In silico cloning study indicated that the designed vaccine candidate might show high-level expression in the bacterial host system. The immune simulation analysis suggests that the vaccine might elicit a strong immune response and when compared to other similar publications which have reported immune simulation for the designed vaccines, our designed vaccine candidate induced a higher level of immunoglobins (IgM + IgG), immunocomplexes and interleukins (IL-2). The Simpson index, D which indicates a measure of diversity, predicted this vaccine candidate to have a diverse immune response. The findings of this in silico studies need to be validated through various immunological assays.

## Acknowledgements

We are grateful to Manipal-Schrödinger Centre for Molecular Simulations, Manipal Academy of Higher Education and Manipal College of Pharmaceutical Sciences for providing necessary supports and facilities to carry out the present research work.

## Abbreviations

SARS-CoV-2: severe acute respiratory syndrome coronavirus 2
ACE-2: angiotensin-converting enzyme 2
HCoVs: human coronaviruses
RBD: receptor binding domain
TLR-3: toll-like receptor.

